# The effect of atmospheric pollution caused by pheasant *Phasianus colchicus* releases on the epiphytic flora on trees in sensitive woodlands

**DOI:** 10.64898/2026.05.08.723433

**Authors:** R.B. Sage, C. Bealey, M.I.A. Woodburn, J. Werling, A.N. Banks, D. Abrahams, J.R Madden

## Abstract

The release and management of pheasants (*Phasianus colchicus*) in the UK for recreational shooting exerts a range of effects on the ecosystem into which they are released. We studied possible effect of nutrient deposition on epiphytic tree flora at 20 pheasant release sites distributed through England (18) and Wales (2) during winter and spring 2023/24. Sites were all Ancient Semi-natural Woodlands (ASNWs) and had substantial (600-8000 pheasants) in a single release pen.

We measured N-sensitive and N-tolerant indicator bryophyte and lichen species on tree trunks near to the pen and then in plots along a transect 100m, 250m, 500m and 1km+ away from the pen. To achieve a gradient of pheasant use, the transects were located in the opposite direction to the game managed / shooting area.

We recorded 1.9 times more coverage of N-tolerant lichens and bryophytes combined on selected tree species at the pen-edge compared to the control plots. The relationship showed a decline from the pen edge to 250m away but then stabilised. We also detected higher levels of coverage of N-sensitive tree flora at 100m and 250 m compared to the penedge plot. These measures were also higher at these mid distances compared to the 500m and 1000m plots. We suggest far plots were nearer wood edges and were affected by ambient inputs of aerial N from farmland and other external sources.

The overall interpretation is that concentrations of pheasants in and around release pens for several months from late summer until early winter in ASNWs does affect the balance of N-sensitive and tolerant tree flora up to potentially 250m and this is a consideration when locating release pens in and near to sensitive woods.

## INTRODUCTION

Common Pheasant *Phasianus colchicus* releasing is undertaken in woodlands throughout the UK to support driven game shooting (Tapper 1992, Madden & Sage 2020, Sage *et al*. 2021). Pheasants are released in a number of other countries including USA, New Zealand, Denmark, France and Italy) for the same purpose but usually at lower numbers/densities (Madden & Sage 2020; Powolny & Czajkowski 2022,). In the UK several tens of millions are released annually (Aebischer 2019, Madden 2021). When large numbers are released, they can produce densities in the landscape on shooting estates of up to two thousand birds per km^2^ in late summer (Sage et al. 2025). Typically, the birds are artificially reared from day old-chicks and in late summer when they are 6-8 weeks old, placed into large woodland-based open-topped pens in woodland to protect them from ground predators (e.g. red foxes *Vulpes vulpes*) as they acclimate to independent living (GCT 1996). The basic aims of the management of these releases in and around woodlands is to softly introduce them to the environment, keep birds healthy and protect them from predators, and to provide attractive habitat and feeding points to hold birds in the right places to facilitate driven shooting during the following winter (GCT 1988, 1996, Sage *et al*. 2021).

It is estimated that in the UK, one in 12 woodlands contain a release pen (Sage *et al*. 2005), with at least 14% of woodland in the UK (25% in England) managed in part for gamebirds, primarily pheasants (Gilbert 2007). Consequently, any ecological effects that these releases might have are likely to be of widespread interest. There is interest in the UK about the potential effects these released birds might have on ecologically sensitive areas, particularly those with a conservation designation (Minter *et al*. 2024)

The direct ecological effects of releasing pheasants on woodlands can be split into management effects and the direct effects of the birds themselves. Gamekeepers will manage woodland and adjacent habitats and provide feed and other resources (e.g. shelters) with the broad aim of holding birds at release sites during early life. They then encourage them to move to other places usually outside of the woodland, to facilitate driven shooting (GCT 1996). This management work can improve woodland and the adjacent habitats, for some other wildlife groups including most wild birds (see Madden & Sage 2020, Mason *et al*. 2020, Sage *et al*. 2020, 2021, Madden 2023, for a review of those effects).

The direct effects of the pheasants themselves, again very generally, can damage woodland habitats around release pens and have negative local ecological effects (see the same reviews). These can include physical damage through trampling and pecking, leading to crushed vegetation and bare ground (Sage *et al*. 2005, Sage 2018a, Madden *et al*. 2026). Changes to habitats especially a release and feed sites will also affect invertebrate communities (Humphrey *et al*. 2005, Kirby *et al*. 2005, Hall *et al*. 2021, Sage *et al*. 2026). A second form of direct effects is eutrophication of woodland soils and air through the deposition of nutrients in faeces. The nitrogen (N), phosphorous (P) and potassium (K) remaining from the grains or high protein pellets that the birds have been fed may become concentrated in areas of high gamebird occurrence. Anecdotal reports describe local soil enrichments accompanying large gamebird releases (Alsop & Goldberg 2018, Smith 2014). Sage et al. (2005) found that soil K and P levels (not N) were higher inside release pens than nearby control areas. Capstick *et al*. (2019) found that these differences might persist for over a decade. However, Madden *et al*. (2026) found no evidence that nutrient levels (N, K, P) at a range of distances outside release pens differed from those at control sites >1km from any releases, suggesting that the depositions are highly localised.

There is some anecdotal or unpublished evidence that lower order plants (bryophytes and lichens) in pheasant release woodlands may be affected by atmospheric nutrient enrichment caused by those pheasants (Bosenquet 2018, Sage 2018a,b). Lichens obtain almost all their nutrients from the atmosphere through uptake over their entire surface. They have no means of controlling nutrient uptake, unlike vascular plants, and free exchange of both gases and solutions occurs across cell surfaces (Turetsky 2003). In addition, their surface area to mass ratio is very high and capacity for assimilation is relatively low and therefore pollutants tend to accumulate in the thallus. Lichens are therefore highly susceptible to changes in atmospheric chemistry and deposition and for this reason can be used as sensitive indicators of such changes.

In Europe lichens have been used as sensitive bioindicators of air quality for more than a century and continue to be used for similar research (Sebald *et al*. 2022). Not all lichens are sensitive to any particular pollutant, while some can be highly tolerant. Species are variously sensitive to sulphur, nitrogen, acidity, halogens (e.g., fluoride), heavy metals and ozone (Van Dobben *et al*. 2001). Long distance transport of nitrogenous air pollution is an important driver influencing the occurrence of acidophytic (acid loving) lichen species, which are usually the most important in conservation terms, and is a real threat to these populations (Van Herk 1999). Reasons for sensitivity to N compounds in acidophytic lichens include an increase in bark pH i.e., lower acidity (often caused by ammonia), effects of NH_4_ ^+^and or NO_3_^-^ in precipitation (N-deposition) and eutrophication causing increased and dominant growth of competing species such as algae (Mitchell *et al*. 2005). In the past, acid rain was put forward as the main cause of the decline in ‘acid sensitive’ *Lobarion communities* (Gilbert 1986). Changes to bark pH however may not be the main driver of species composition change which may also be attributed to nitrogen (Farmer *et al*. 1992). Some evidence suggests that the concentration of N, particularly as ammonia, may be more important than the overall N dose (Britton & Fisher 2010).

More recent research (Seed *et al*. 2013) has identified species of lichens on oak and birch trees (*Betula* spp.) across the UK that are sensitive to, or tolerant of, increasing concentrations of gaseous nitrogenous pollutants. This information has been used to formulate a simplified suite of species for monitoring air-borne pollution in the UK. The response to increasing atmospheric N pollution in the field can be measured by the decrease in N-sensitive and increase in N-tolerant lichen species growing on the bark of these tree species (Wolseley *et al*. 2004).

Bryophytes (mosses and liverworts) are among the most sensitive of vegetation taxa with respect to pollutant deposition and can be sensitive to both acidity and N levels, both of which are the most dominant elements of anthropogenic deposition, resulting in both N-sensitive and N-tolerant mosses. Too much N can, for example, change morphology, leading to sparser mats that are sensitive to desiccation and less efficient at suppressing competitors (Armitage *et al*. 2012) and photosynthetic processes can also be compromised. A European survey (Harmens *et al*. 2011) found that the N concentration in two common mosses correlated well with modelled N deposition. They suggested that these and other mosses could be used to provide reliable estimates of nitrogen deposition, i.e., indicator species. It has also been suggested that high nitrogen pollution can lead to excessive algal growth, possibly smothering bryophytes on trees (Wolseley *et al*. 2004). Hill et al. (2007) produced ‘attribute data’ for all British and Irish mosses and liverwort plant species. This includes assigning them Ellenberg values (Ellenberg, 1974), on a nine-point scale indicating their tolerance to background environment including nitrogen (based more on soil fertility or productivity, not presence of mineral nitrogen). This has enabled ecologists to interpret distribution patterns in response to environmental pressures. This was updated in 2017 (<u>BRYOATT - Attributes of British and Irish Mosses, Liverworts and Hornworts - Spreadsheet | Biological Records Centre</u>).

For this study, we recorded a number of selected N-tolerant and N-sensitive bryophyte species, together with the set of lichens, to explore any changes to the tree-flora community that may be attributable to pheasant feeding and defecation leading to atmospheric N-enrichment. We recorded the tree species that they grew on to account for any differential effects of bark accumulation of nutrients. We extended the surveys beyond the immediate release site to account for dispersal of the pheasants and their reduced densities. We used sites >1km from any releases as controls. Our expectation was that responses to any effect of N-enrichment would vary with distance from the release pen, with fewer measurable effects further from the main point source of pollution (i.e. the release pen).

## METHODS

### Sites

We collected data at 20 study sites in England (18) and Wales (2), in areas of ancient semi-natural broadleaved woodland. The strategy was to sample tree flora on several trees in each of four plots located at increasing distances up to 500m from a pheasant release pen in the same wood, and then in a further control plot at 1km or more from the release pen. This 500m requirement defined a minimum woodland size for consideration and was stipulated by the current licencing requirements for gamebirds in the UK in which releases on some protected areas either require consent (SSSI) or if within a 500m buffer zone for SACs and SPAs, a licence, the type of which varies depending on type of protected area and the conditions of release (https://www.gov.uk/guidance/gamebirds-licences-to-release-them). Fieldwork was conducted throughout the winter of 2023-4 and into the early spring of 2024 and included sites across 12 counties in England and Wales to attempt a representative sample of woodlands, with a focus on regions where large-scale releases and shooting is most common (Table 1).

**Table 1.** The distribution of the twenty ASNW sites used in the study by county across England and Wales. Locations are given at a county level to ensure shoot anonymity.

| Country/County name | Number of Sites |
| --- | --- |
| <u>England</u> |  |
| Cambridgeshire | 1 |
| Cornwall | 2 |
| Cumbria | 1 |
| Devon | 2 |
| Hampshire | 3 |
| Herefordshire | 1 |
| Kent | 2 |
| Leicestershire | 1 |
| Northamptonshire | 1 |
| Somerset | 1 |
| Staffordshire | 1 |
| Wiltshire | 2 |
| <u>Wales</u> |  |
| Denbighshire | 1 |
| Powys | 1 |

All pens in potential woodland sites needed to have been releasing gamebirds each year for the past five years. Initially we asked sites if the release into the study pen was upwards of 800 pheasants. Subsequently we asked for further details about the pen release. Not all sites disclosed the actual release number, but following our site visits we were confident that all 20 sites released between 600 and approximately 8000 pheasants into the focal pen. Study sites also needed to include suitable woodland for a control plot at least 1km from the pen. At some sites this plot was in the same woodland block as the rest of the transect, while at others it was in a separate block containing a similar tree community. All control plots were between circa 1km and 3 km from the main study pen. An additional consideration was the presence of other significant nitrogen enrichment sources. Sites that were close to busy roads, large-scale pig units or poultry farms or other potential sources of aerial pollution, were excluded. Realistically, it was impossible to avoid all possible sources of background pollutants that could potentially influence our results.

### Transect location

In woodlands, released pheasants usually make daily movements between release pens and sites where they are deliberately attracted by gamekeepers using supplementary feeders, game crop plots and other specific game management habitat where the resources they are looking for have been located or created (GCT 1988; GCT 1996).

Our aim was to investigate potential effects of released pheasants on tree flora on trees adjacent to release areas and then along a transect moving away from that area, while deliberately avoiding feed points or other places where pheasants were encouraged to congregate. These congregation points are directly influenced by game management, so in the case of sensitive areas, game managers could deliberately move dispersing birds away from such areas and instead concentrate them in less sensitive areas. Thus, we expect direct ecological effects to be high at such congregational sites, regardless of their distance from the release pen. Instead, we looked at areas where birds were moving without directed management, which could include intrusions into sensitive areas where unintended congregations and hence damage might occur. If the dispersing birds were causing these direct effects, then we would expect to detect a gradient of effects that declines with distance from the release pen. It was therefore necessary to have an understanding of game management activities on study sites so as to exclude these attractive, managed areas from our transects and sampling locations. Sampling plots were always located further away from other game management interests than the study pen. Therefore, we selected a transect that led from the release pen in as direct a line as possible, while remaining within the woodland block, and avoiding management sites and any other point sources of nutrient pollution. Pheasant counts were undertaken at all sampling plots and there was a clear declining trend of decreasing encounters with distance (reported in Madden *et al*. 2026).

Having defined a suitable transect orientation, we located sampling plots as follows: within 5-10m of the release pen edge and then at 100m, 250m, and 500m from the pen plus a control plot circa 1km+ away. The position of each sampling plot was recorded using a hand-held GPS device (level of accuracy approximately 5-10m, depending on canopy cover, and this introduces some noise in distance measurements. Sampling plot locations along transects had to take account of site considerations, such as woodland type/ tree species. For example, a site where an area of trees might have been cleared in recent times at, say the 250m location but where mature stands existed at 300m, then the sampling point would have been taken at the 300m point. At several sites transects were not in straight lines but straight-line distances between plots were estimated to ensure distance requirements were still approximately equivalent.

A potential further consideration was to take account of prevailing wind direction because it might be expected for atmospheric eutrophication to travel further downwind of a source. In the UK, the prevailing wind direction in September, when pheasants start to disperse from their pens, is from the South-West (Lapworth & McGregor 2006). However, we could not ensure that all transects fell on the same orientation given other requirements described above. Therefore, we post-hoc tested whether patterns of tree flora cover and change in the cover was related to the transect orientation (see below).

### Tree flora assessments

Tree flora assessments were undertaken at all sites during winter and early spring 2023-2024. Winter is an optimum period to conduct surveys of most bryophyte and lichen species as the high atmospheric moisture content means that the bodies of the plants are fully hydrated and therefore relatively easy to identify, despite the fact that many will not have fruiting bodies present https://britishlichensociety.org.uk/learning/studying-lichens.

The first of five trees in each sample point was randomly selected as one meeting the species type and minimum diameter criteria (see details below). The other four trees at each sample point were selected by walking 15 paces from the central tree in each of the four compass/ cardinal directions and using the nearest suitable tree (species & size) to survey.

For each selected tree, the Air Pollution Information System (APIS) - *Monitoring Air Quality Using Lichens* – ‘field guide and app’ method was employed. First, pre-defined tree species only were identified (preferably oak (*Quercus robur*/*Q. petraea*) and birch (*Betula pendula*/*B. pubescens*) but if these were not present, rowan (*Sorbus aucuparia*), alder (*Alnus glutinosa*), beech (*Fagus sylvatica*), lime (*Tilia* spp.) or ash (*Fraxinus excelsior*) These are all relatively acid-barked trees (tree bark in order of reducing acidity: birch, oak (pH 3.8-5.8), rowan, alder, beech, lime, ash (pH 5.2-6.6). Second, odd trees growing in for example dense shaded clumps were avoided, as were trees with a high cover of Ivy (*Hedera helix*) i.e., where it largely obscured the tree trunk and made locating quadrats difficult. Wherever possible, selected trees were single stemmed (standard) with a straight trunk, > 40 cm in circumference). Third, the three aspects (East, South and West) on the trunks to be surveyed were selected using a hand-held compass and a 50 x 10cm quadrat on each of the three aspects of the tree between 1.0 and 1.5 m above ground level was delineated (Wolseley et al. 2004). Fourth, presence and % cover of N-sensitive and N-tolerant indicator species within each trunk-quadrat were recorded. Species recorded included:

1. Lichens: **N-sensitive indicators** (as listed in APIS): *Pseudevernia furfuracea, Evernia prunastri, Hypogymnia spp, Parmelia spp, Usnea spp, Bryoria spp, Sphaerophorus globosus, Graphis spp, Ochrolechia androgyna;*
2. Lichens: **N-tolerant indicators** (as listed in APIS): *Xanthoria parietina, X. polycarpa/X. ucrainica, Candelariella reflexa, Punctelia subrudecta, Physcia adscendens/P. tenella, Anthonia radiate, Lecidella elaeochroma, Amandinea punctata*.
3. Bryophytes: **N-sensitive indicators** (Ellenberg N indicator values of 2): *Dicranum scoparium, Frullania tamarisci;*
4. Bryophytes: **N-tolerant indicators** (Ellenberg N indicator values of 6): *Amblystegium serpens, Brachythecium rutabulum, Rhynchostegium confertum, Thamnobryum alopecurum*.

The four highly N-tolerant and two highly N-sensitive bryophyte species selected were deemed to be sufficiently widely distributed and hence likely to occur across the survey area. They were selected in consultation with the project team at Natural England and added to the standard list of lichens to be recorded.

## Data analysis

### General

Analysis of the tree flora variables was aimed at identifying distance trends by comparing values from trees in each plot with other plots in the transect, and with the control plot. Two models were constructed, one considering cover by N-sensitive tree flora that we expected might be low close to the pen where airborne pollution levels were high, and increase further from the pen as air-borne nutrients from released pheasants declined, and the second with N-tolerant tree flora that we expected to show the opposite pattern.

The dependant variable for analysis was constructed from the sum of cover (with cover <1% rounded to 1%) for each of the N tolerant or N sensitive species groups. Lichens and Bryophytes were combined. Analyses were conducted using generalised linear mixed models to account for the use of repeated transect measures across multiple study sites. A generalised model structure was used because dependent variables were proportions with negative binomial functions and log links. The general structure was to include the five ‘distance-from-the-pen’ categories as the main independent variable of interest with the Tree Species included as an additional variables to account for any species effects, and the interaction between Tree Species and Cover to account for any differential effects of the tree species affecting the tree-flora cover. We included site as a random term to allow patterns to differ between sites that may be expected due to underlying environmental differences at the sites which we had not accounted for. All analyses of effects of distance on cover were conducted using the r package *glmmTMB* (Brooks *et al*., 2017) in R (R Core Team, 2024).

This allowed us to ask two questions in each test. First, did the measure of tree flora just outside the pen, circa 5-10m away (Plot A) differ from that collected at the control site (Plot E), at least 1km away from the pen i.e. in a woodland area where no gamebird management was conducted? Second, at what distance (100m, 250m, 500m, 1000m) could we detect any differences from the measure collected just outside the release pen? We addressed these questions by conducting post-hoc contrast tests using the *emmeans* package (Lenth, 2025) with Tukey correction for multiple comparisons. Probability values of main effects or interactions were derived using the “drop1” function. We inspected model fit visually. We checked measures of dispersion and zero inflation using the *DHARMa* package (Hartig, 2022). Models were not over-dispersed (all p > 0.33) or zero-inflated (all p > 0.91).

### Testing the effect of wind direction

We tested whether there were differences in the area of cover, or the rate of change of cover with distance, for both N tolerant and N sensitive tree flora, depending on the orientation of the transect relative to the prevailing wind direction. We calculated what was the transect angle where peak area of cover or rate of change of cover occurred. We then tested whether this peak orientation, and declines around it, correlated with the assumed prevailing wind direction around and after the time of release, from the south west (225°) (Lapworth & McGregor 2006). Circular analyses were conducted using the *circular* package (Agostinelli and Lund 2025).

## Results

### Nitrogen Tolerant Indicator Species

Quadrat cover of nitrogen-tolerant indicator tree flora varied depending on the tree species being studied (χ^2^ < _4_ = 35.9, P < 0.001, Figure 1), with lower cover on Beech and Oak compared to Ash, Birch et al. (post-hoc contrasts, Beech vs Ash, Birch *et al*., all p < 0.007, Oak vs Ash, Birch *et al*., all p < 0.055).

**Figure 1.**
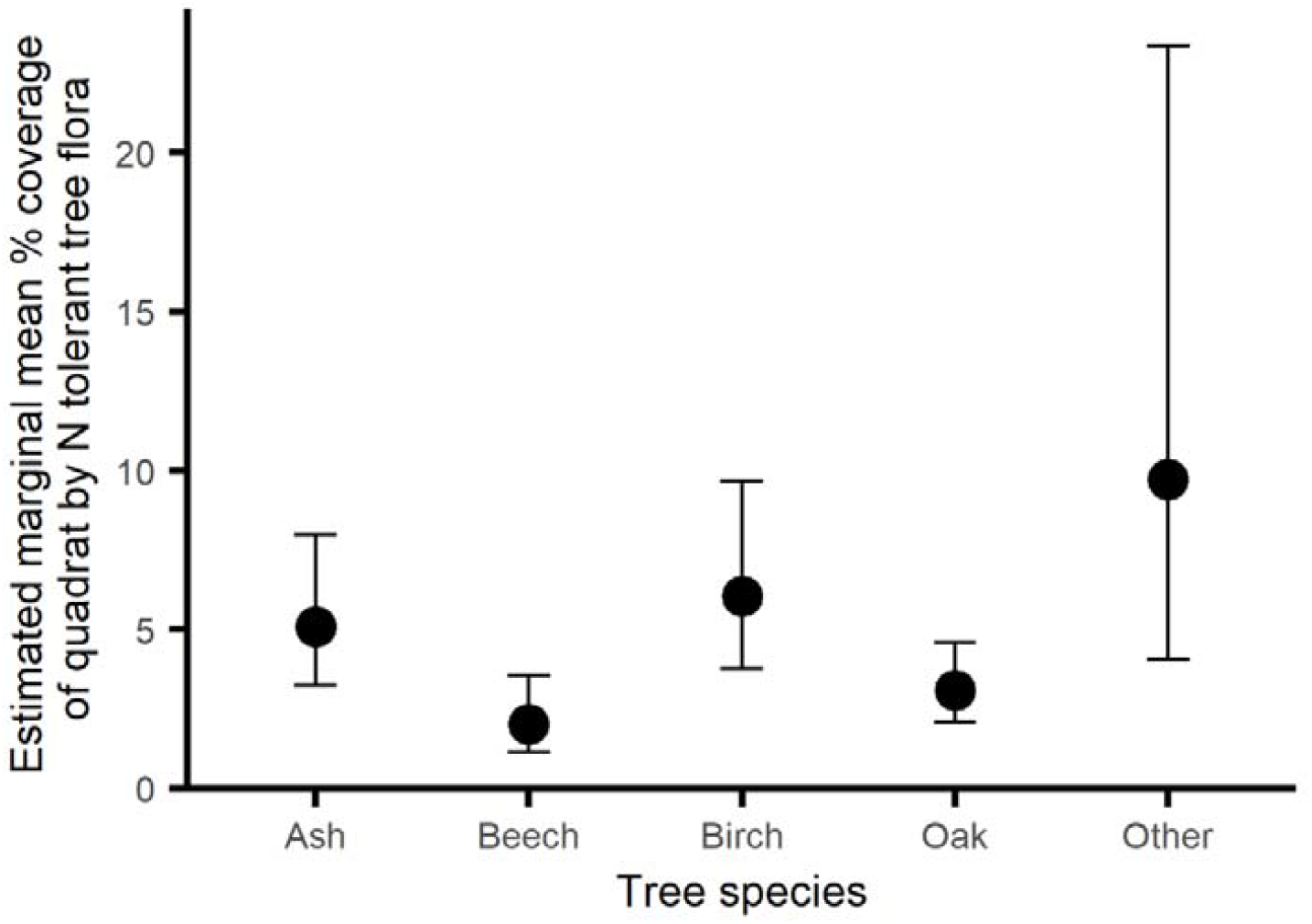
Percentage cover by N-tolerant tree flora (lichens and bryophytes) in quadrats on trees of different species during winter/ early spring surveys. Error bars indicate 95% CI.

While including tree species in the model, the quadrat cover by nitrogen-tolerant indicator tree flora differed depending on the distance from the release pen (χ^2^ _4_ = 20.2, p = 0.0058, Figure 2). There tended to be 1.9 times greater cover of N-tolerant epiphytic tree flora in quadrats just outside the release pen compared to the control sites (post-hoc contrast, p = 0.096). There was significantly lower cover compared to just outside the pen at 100m and 250m (post-hoc contrasts, 100m p = 0.0017, 250m p = 0.015).

**Figure 2.**
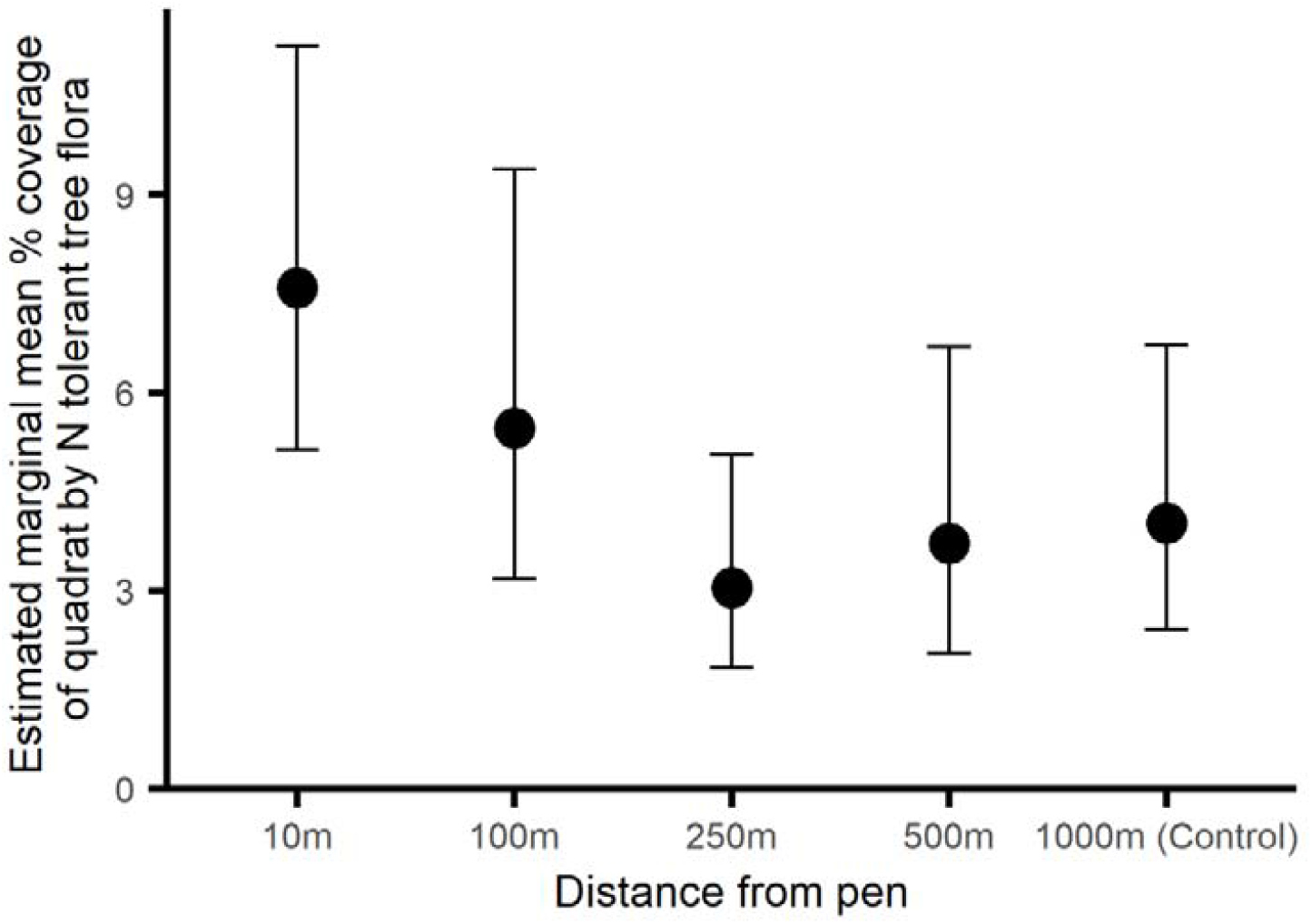
Percentage cover by N-tolerant tree flora (lichens and bryophytes) in quadrats on trees at different distances from 20 woodland pheasant release pens. Error bars indicate 95% CI.

### Nitrogen Sensitive Indicator Species

Unlike the N tolerant indicator species, quadrat cover by nitrogen-sensitive indicator tree flora did not vary depending on the tree species being studied (χ^2^ _4_ = 5.31, p = 0.25).

However, because Tree Species had been influential for N tolerant species, and because previous work indicates that different barks may accumulate nutrient differentially (see Introduction), we retained Tree Species in our model. The nitrogen-sensitive indicator tree flora cover differed depending on the distance from the release pen (χ ^2^ _4_ = 23.1, p = 0.0014, Figure 3). Cover just outside the pen did not differ from that at control sites (post-hoc contrast, p = 0.93). However, there was significantly higher cover compared to just outside the pen at 100m (post-hoc contrasts, 100m p = 0.0031).

**Figure 3.**
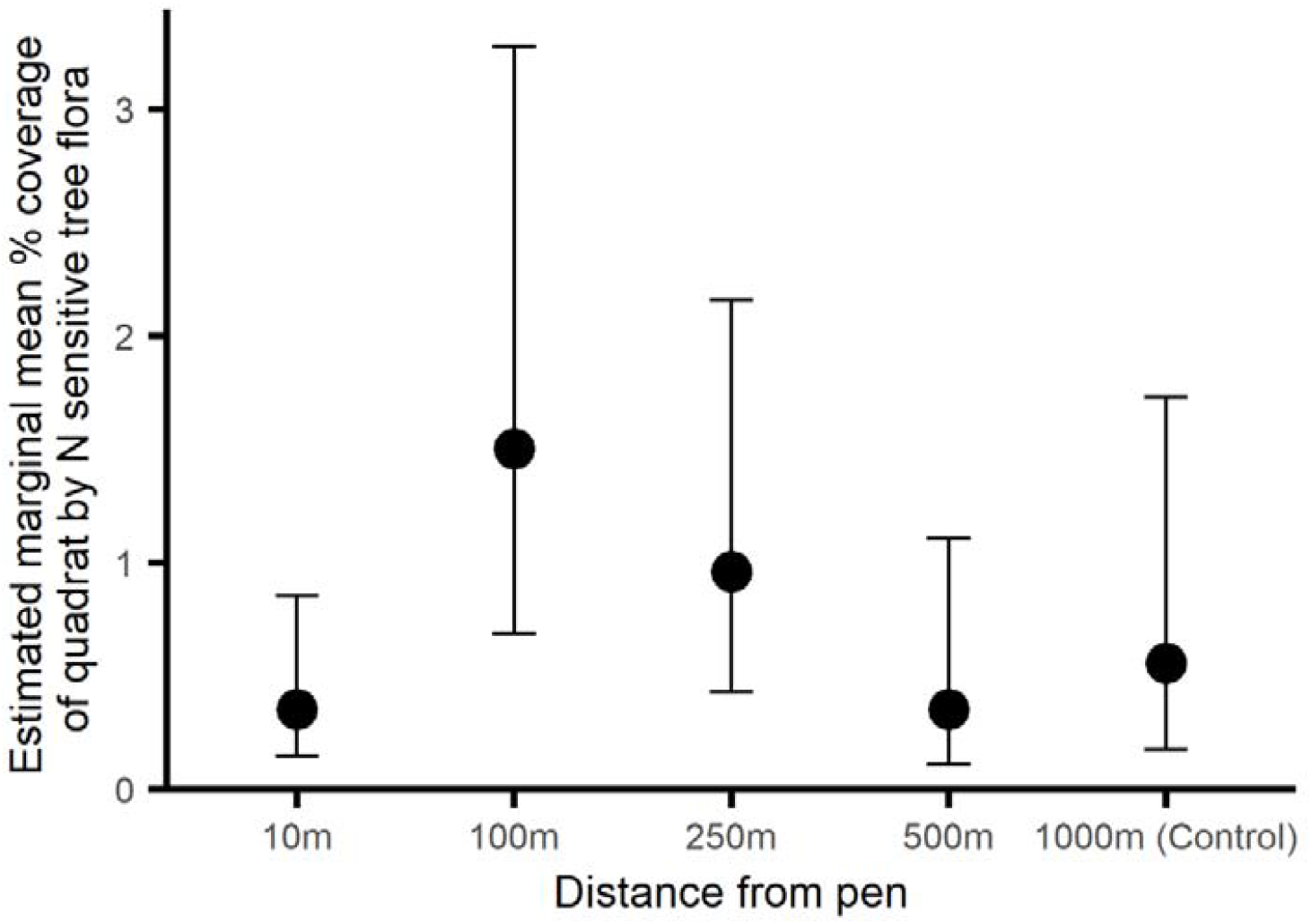
Percentage cover by N-sensitive tree flora (lichens and bryophytes) in quadrats on trees at different distances from 20 woodland pheasant release pens. Error bars indicate 95% CI.

### Effects of Transect Orientation

We found no evidence that either the total cover by N-tolerant or N-sensitive tree flora, or the strength of change in cover with distance for either group, was related to the prevailing wind (South West, 225°) (Table 2).

**Table 2.** The relationships between the deviation of the transect orientation from the prevailing wind direction (225°) and the overall N-sensitive or N-tolerant tree flora cover, or the change in that cover with distance from the central pheasant release pen.

|  | Peak orientation<br>(°) | Relationship between the deviation of the<br>transect orientation from the prevailing wind<br>direction (225°) and the tree flora cover<br>measure |  |  |  |
| --- | --- | --- | --- | --- | --- |
|  |  | Est | SE | t | p |
| Overall cover by N tolerant tree<br>flora (N tolerant intercept) | 161 | 1.41 | 4.57 | 0.31 | 0.76 |
| Change in cover with distance<br>from the pen by N tolerant tree<br>flora (N tolerant slope) | 9 | -0.74 | 0.91 | -0.81 | 0.43 |
| Overall cover by N sensitive tree<br>flora (N sensitive intercept) | 31 | -1.60 | 1.39 | -1.15 | 0.26 |
| Change in cover with distance<br>from the pen by N sensitive tree<br>flora (N sensitive slope) | 224 | 0.69 | 0.40 | 1.73 | 0.10 |

## DISCUSSION

We found that trees exhibited higher cover by a suite of N-tolerant tree flora (both lichens and bryophytes) close to woodland pheasant release pens. This cover declined further from the pens such that cover tended to be almost twice as high close to pens as at the 250m and 500m plots, and at the control sites >1000m from any release sites. Additionally, tree cover by a suite of N sensitive tree flora was lower close to the release pen compared to cover 100m away, but declined at further distances, such that cover at control sites did not differ from that at the release pen. As expected, we found that cover differed according to the tree species surveyed. We found no evidence that patterns of either overall cover or change along the transects from pen to control sites varied depending on the transect orientation relative to the national prevailing wind direction.

Our interpretation of the N-sensitive tree flora result is that in plots immediately next to the release pen, hosting higher estimated pheasant activity (Madden *et al*. 2026), the sensitive tree flora were suppressed by aerial pollution from pheasants in the pen compared to plots 100 and 250 m away. Cover was also supressed in the 500 m plot and the control, but here we suggest aerial pollution from sources outside the wood. Our interpretation of the N-tolerant tree flora result is that in plots close to the pen, the higher levels of aerial nitrogen either directly stimulated higher growth of N-tolerant flora or it suppressed N-sensitive flora, enabling the tolerant species to thrive. The decline in cover by N-tolerant flora out to 250m suggests a decline in levels of aerial nitrogen originating from the high concentration of released birds inside and close to the pens. The slight increase in cover at greater distances, out to the control plots suggests that alternative or additional sources of airborne nitrogen may also influence tree flora cover. Therefore, our results suggest that airborne nutrients, specifically nitrogen in all forms, originating from released pheasants, might affect lichen and bryophyte community composition on trees up to 250m from woodland release pens.

This result can be compared with that from Sage (2018a,b) that looked at sites in the SW of England and suggested that the abundance and diversity of bryophytes and lichens on trees overall was not different between release pen plots and control plots away from pens in the same woods or in woodland without pheasant release pens. Our focus on known N-tolerant and N-sensitive flora may explain why we found distance effects compared to the broader abundance and diversity measures used in the previous studies. Similarly our consideration of a finer-scale set of distance bands also revealed subtle changes over shorter distances than those considered previously However, moss diversity was reported to be about 25% lower on trees in both release pen plots and in the control plots outside the release pens but in the same woods in Sage (2018a,b), compared to the nearby estate control woods without release pens. Liverwort species diversity was between approximately 30-50% lower in the release pens and release pen wood controls, compared to the non-pen estate control woods. Taken together, these earlier results provide an indication that large releases in woodland pens can affect lower order plant communities on trees away from the release pens but in the same wood and hence supports the more focussed (in terms of distance) findings of the study reported here.

We can also compare our results with similar studies investigating effects of air-borne pollution from agricultural sources. Pitcairn *et al*. (1998) report four field studies measuring ammonia along transects downwind at two poultry farms, a pig farm and a dairy farm. The study documents ammonia concentrations moving away from the farms which return to background levels well within 1 km from the source. In particular, changes in key flora metrics with distance from the poultry farms (holding between 100,000 and 200,000 broilers) in woodland were limited to within 300m or so. Pheasant releasing is undertaken at a relatively small scale compared to poultry farming, and releases at our sites did not exceed 6000 birds, so represented almost two orders of magnitude less than the poultry sources studied.

One potential alternative explanation for the distance effects we report is that release pens are usually located at or close to wood edges. However, because of our aim to avoid game managed areas in our study (apart from the release pen itself) we were invariably working from the side of pens set deepest into the woodland, with transects from those starting points taking us initially at least, deeper into the main woodland. Therefore, the pen plot was exposed to aerial pollution from the pen, but actually protected from other external sources of pollution outside of the wood, such as ploughed fields, roads and livestock. However, at the far end of the transect at 250 – 1,000m we were frequently approaching wood edges and hence greater exposure to these external pollution sources than both the pen edge plot and the 100 and 250 m plots.

Our results exhibited high levels of variation between sites. Despite our efforts to ensure standardised sampling conditions, we were restricted by local geographical and game management factors, and these additional variables that we did not include in our analyses due to having a sample of only 20 sites may explain some of this variation. One source of variation is that differences in canopy cover, either due to woodland composition, age or structure may explain differences in aerial N that can penetrate from prevailing winds, via inputs from farming or other land use. This was probably most influential at the 500m and control locations. Both a reduction in overall age class (Király *et al*. 2013) as well as wider landscape effects such as regional deciduous tree cover (Möller *et al*. 2025) can also affect tree flora cover. Another factor determining the relative abundance of N-sensitive and N-tolerant lichens and bryophytes at sites could be how much biomass occurred on the trunks of the trees surveyed. Trees in the wetter western woods can host a high biomass of bryophytes (mainly mosses) and this could increase the amount of airborne pollutants held among the plants and therefore filtering down to the bark (Mitchel *et al*. 2004). It is possible that this results in a reduced sensitivity to further eutrophic factors such as presence of a pheasant release pen. Also, the high biomass of mosses may out-compete or obscure the lichens. A final key potential factor is local instances of intensive animal husbandry which are a major contribution to nitrogen deposition on the ground in the UK and elsewhere and which are shown to affect depositions away from sources (Dragosits *et al*. 2006). We did not have information about farming activities outside our study sites and so although we avoided such known point sources, they may occur off-site but within range to affect airborne nutrient levels. Dragosits *et al*. (2006) summarise data from Pitcairn *et al*. (1998) and other sources collected in the late 1990s-early 2000s, and conclude ammonia concentrations downwind of large sources declined to (upwind) background values within 1 km. The research indicates, without giving figures, that the size of the source affects the distance over which concentrations are increased.

Two sources of variation that we did expect and consider in our analyses were the tree species hosting the flora and prevailing wind direction. The substrate tree species explained significant variation in cover by N-tolerant but not N-sensitive species. We included the species in our models to control for this variation. Contrary to our expectations, we found no evidence that transects that ran downwind from release pens either had lower cover by N-sensitive flora or exhibited shallower patterns of change as distance increased. This might be because although the prevailing wind in the UK in autumn is from the south west, there is sufficient variation over time to obscure any patterns, or that the woodland density and transect orientation interacted such that wind flow was differentially disrupted across sites, again obscuring any patterns.

While there are release pen stocking density restrictions at or within 500m of designated sites in England (not Wales) as a part of current licencing arrangements (Defra 2024), there are not pen size restrictions, so numbers released at designated sites included in our study are not restricted and hence in the context of our study not necessarily lower than at other sites.

Our results, considered alongside previous work on pheasant releases in Exmoor (Sage 2018a,b) and effects of large-scale poultry farming (Pitcairn *et al*. 1998), suggest that typical pheasant release pens containing a few thousand birds or fewer, who are concentrated there for a relatively short time, may have an effect on the cover and community composition of N-tolerant and N-sensitive tree lichens and bryophytes at sites within a few hundred metres of the release pen. At distances beyond this, other external factors, may have a bigger impact. This suggests that the releases of pheasants should be carefully evaluated if situated less than 250m from protected areas containing populations of sensitive tree flora.

## Acknowledgements

We thank all the estate owners for granting us permission to carry out the surveys on their estates, NE site managers for facilitating access to designated sites and the gamekeepers for their help at all sites. We thank Simon Curson, Jonathan Cox and Matt Wainhouse at NE for advice on fieldwork protocols and data variables; and Nicholas Aebischer at GWCT for statistical advice, especially during the early design stage. Thanks to staff at Defra for providing funding and oversight, especially David Rymer. Natural England commissioned this research as part of Defra’s Gamebird Research Programme. Natural England contracts a range of reports from external partners to provide evidence and advice to assist us in delivering our duties. The views in this paper do not necessarily represent those of Natural England or Defra and should not be taken as specific advice.

